# SecDATA: Secure Data Access and de novo Transcript Assembly protocol - To meet the challenge of reliable NGS data analysis

**DOI:** 10.1101/2023.10.26.564229

**Authors:** Sudip Mondal, Namrata Bhattacharya, Troyee Das, Zhumur Ghosh, Sunirmal Khatua

## Abstract

Recent developments in sequencing technologies have created new opportunities to generate high-throughput biological data at an affordable price. Such high-throughput data needs immense computational resources for performing transcript assembly. Further, a high-end storage facility is needed to store the analyzed data and raw data. Here comes the need for centralized repositories to store such mountains of raw and analyzed data. Hence, it is of utmost importance to ensure data privacy for storing the data while performing transcript assembly. In this paper, we have developed a protocol named *SecDATA* which performs de novo transcript assembly ensuring data security. It consists of two modules. The first module deals with a framework for secured access and storage of data. The novelty of the first module lies in the employment of distributed ledger technology for data storage that ensures the privacy of the data. The second module deals with the development of an optimized graph-based method for de novo transcript assembly. We have compared our results with the state-of-art method de Bruijn graph and the popular pipeline Trinity, for transcript reconstruction, and our protocol outperforms them.

## 1. Introduction

NGS technology has facilitated transcriptomic studies through RNA-seq data analysis [1]. To get the complete picture of a biological system, it is important to analyze such high-throughput data efficiently. Such a process includes assembling short reads which helps in the reconstruction of the full-length transcripts. It helps in detecting novel transcripts and identifying alternatively spliced genes as well. Various transcript assembly algorithms [2] have facilitated such analysis, to recreate the entire transcriptome even without a reference genome. Nevertheless, the presence of sequencing errors, polymorphism, sequence repeats, etc makes the task of the transcript assembly process often more challenging. Further, identifying the full set of transcripts which includes novel transcripts corresponding to unannotated genes, splicing isoforms, and gene-fusion transcripts [3] makes it more critical.

A significant amount of work has been done regarding the development of various reference-based and de novo transcriptome assemblers [4]. The reference-based method involves transcriptome analysis, primarily by mapping onto a reference genome [5]. On the other hand, the de novo assembly method is primarily used because many non-model species lack a reference genome for transcriptome analysis [6]. Examples of widely used de novo assemblers are SOAPdenovo-Trans, rnaSPAdes, BinPacker, Bridger, Trinity, IDBA-Tran, ABySS, and Oases [7]. De novo transcriptome reconstruction from short reads remains an open problem, due to the lack of comprehensive knowledge needed to reconstruct the transcriptome as completely as possible. It gets even more complicated as the transcriptome varies with variations in cell type, environmental conditions, etc. Ideally, a transcriptome assembler should be capable of detecting full-length transcripts even for lowly expressed transcripts. Two basic algorithms that are generally used for short read de novo assemblers are overlap graphs and de Bruijn graphs [8]. Mira, Phusion, and Newbler are a few of the widely used overlap graph-based assemblers [9]. Overlap graphs compute the overlaps between each pair of reads making this algorithm computationally more intensive than de Bruijn graphs [10]. Hence, most of the de novo assemblers for RNAseq data analysis use de Bruijn graphs generated [11] from the overlap between k-mer decompositions of the reads in the RNAseq data and not between the actual reads. ABySS, Velvet, Oases, and Trinity are popular de Bruijn graph-based assemblers [8]. In the de Bruijn graph [12], a node is represented by a sequence of k nucleotides of a specific length (’k-mer,’ with k significantly shorter than the read length) and nodes are linked by edges if they overlap perfectly with k-1 nucleotides. This representation enables all possible solutions to be enumerated which comprises linear sequences with k-1 overlap. Each path in the graph represents a potential transcript for the transcript assembly process. The representation of the de Bruijn graph in memory is a computational bottleneck for many assemblers [12, 13]. One of the present-day challenges that remain is the use of de Bruijn graphs for the de novo assembly process. This is because a massive computational resource is needed for implementing such a procedure for de novo assembly (i) by creating a graph from the raw sequence data and (ii) subsequent generation of millions of possible paths from these graphs. Along with transcript assembly, secure data handling is an important area of concern for RNAseq data analysis. It is essential to ensure data security before (securing raw data) and after (securing analyzed data) transcript assembly step in NGS data analysis. Hence, the initial input data should remain unchanged and should not be modified by others without permission. Blockchain technology has achieved notable attention for maintaining data security from the business and corporate sector to the healthcare industry and genomic research [14]. The Blockchain system, also known as distributed ledger technology (DLT) [15] as well as distributed or shared ledger, is a consensus of replicated, shared, and synchronized digital data, geographically scattered over multiple places [16]. The main advantage of DLT lies in the fact that it can be used without requiring a central administrator and also centralized data storage. It is like a distributed database on a peer-to-peer network, that is spread across several nodes (devices) where each node replicates and saves an identical copy of the ledger and updates itself [17]. Cryptographic keys and signatures are used for the security of the nodes [18]. DLT handles and shares information across a network of verified users. To support a wide range of decentralized applications, Hyperledger and Ethereum focus on extended processing capability for building distributed ledger networks [19, 20]. However, DLT was not designed to handle large omics-sized data sets [21].

In this paper, we have designed an optimized pipeline *SecDATA* for de novo transcript assembly that adopts a Blockchain-based strategy. The major focus here lies towards implementing (a) a pipeline that accesses secured data with the help of DLT and (b) performs de novo transcript sequence reconstruction from RNA-seq data.

## Materials and Methods

We have two major focuses in this paper: (i) data security and (ii) de novo Transcript assembly. The computational framework of *SecDATA* is shown in figure 1.

**Figure 1.**
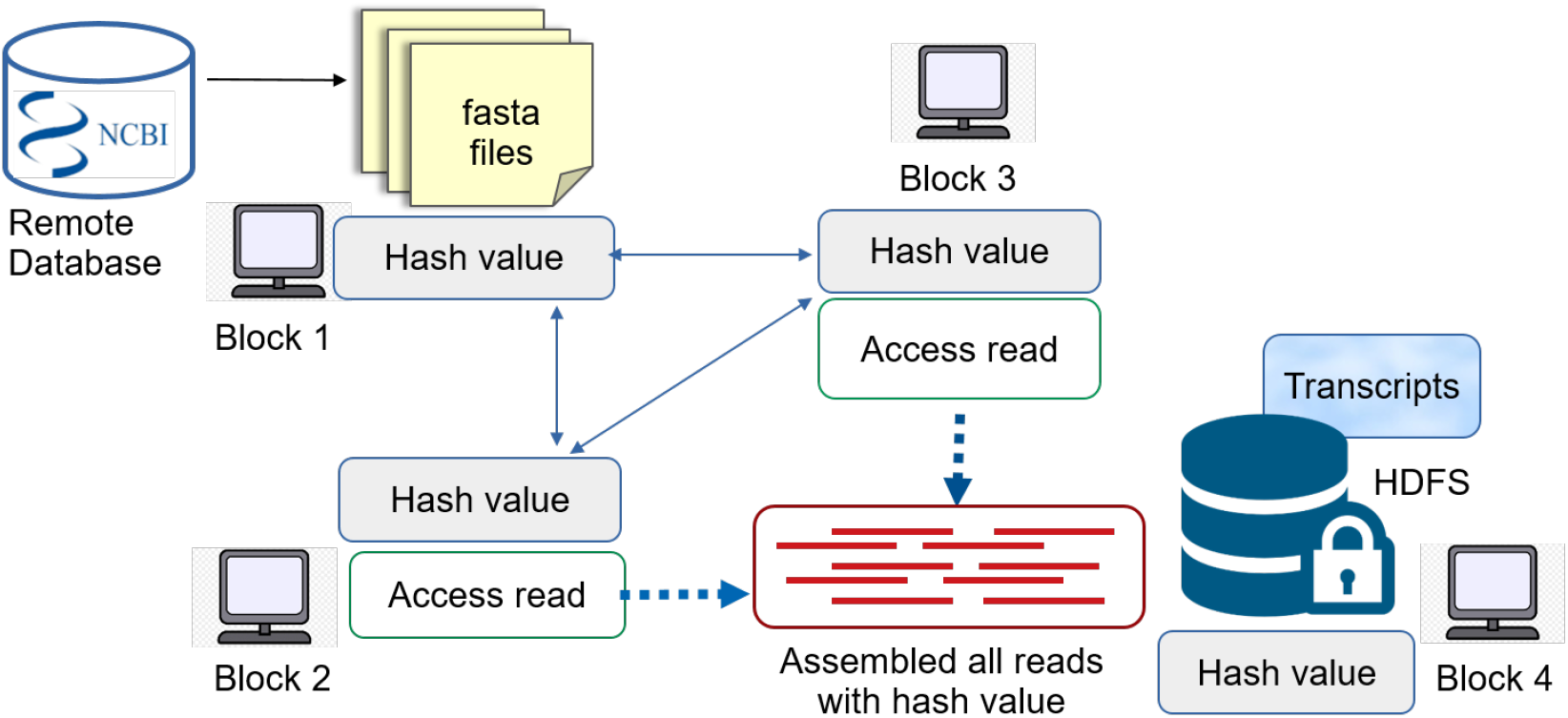
Computational framework for SecDATA

### 2.1. Input dataset and it’s pre-processing

Real and Simulated Datasets: To compare the performance of our transcript assembly protocol, *SecDATA* we have used simulated datasets and publicly available RNA-Seq datasets (dataset provided in Table 1) with different read lengths, number of reads, and depth of coverage. The publicly available data is downloaded from the NCBI (https://www.ncbi.nlm.nih.gov/) SRA database using the streaming technique, ParStream-Seq [22]. We have used Trimmomatic [23] and FastQC [24] for adapter trimming and filtering poor-quality reads and poor-quality bases respectively from the downloaded datasets. We have used a simulation method [25] to generate simulated Illumina reads from a reference genome (human(hg19) chromosome 1 and 2 (Gencode version 18) as reference). The simulated datasets were assembled without any additional preprocessing. Our dataset read length is 100 bp and insert lengths range between 200bp to 350 bp to simulate sequencing reads. The simulated dataset is available at https://github.com/sudipmondalcse/SecDATA

**Table 1:**
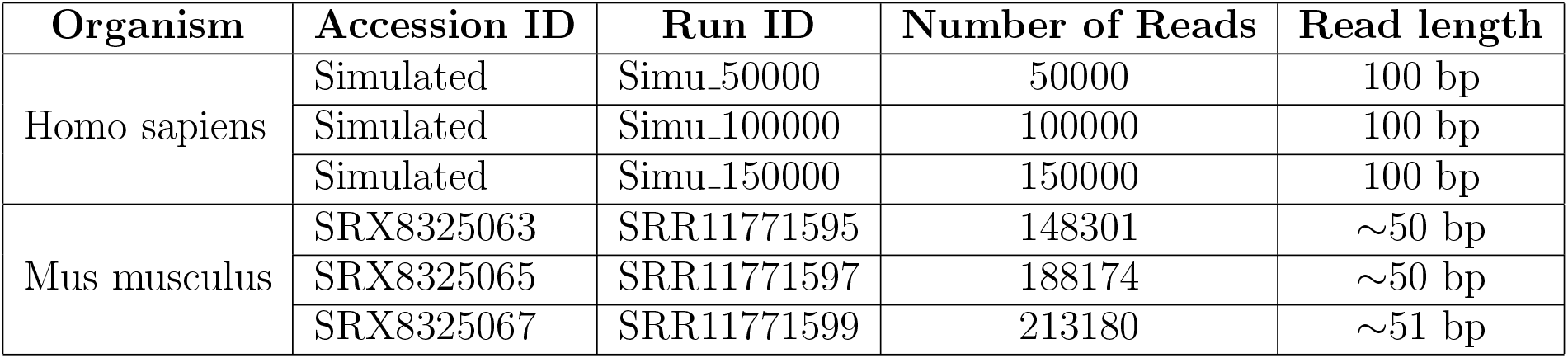
Input dataset.

## 3. Computational architecture of SecDATA

### 3.1. Proposed Model for data security

In this paper, we have used public blockchain technology for secure data storage. A public blockchain is a type of blockchain that is designed to be fully decentralized [26]. Once a record has been added to the chain it is very difficult to change it without permission [27, 28]. Unlike traditional databases, the data stored using this technology is not centralized at one location; instead, the record of the ledger is shared across a network of computers or computing nodes. The network uses an encrypted code to perform constant checking to ensure that all the copies of the database are properly duplicated in all nodes and ensure the privacy of their data [29]. A consensus algorithm [30] achieves data reliability in the Blockchain network and establishes trust between unknown peers by allowing them to reach a mutual agreement about the current state of the distributed ledger. Through a consensus mechanism, all active network nodes ensure that modifications in the database in the form of digital records, follow a set of rules, making the DLT tamper-proof. Many forms of DLTs are available, such as Blockchain like Bitcoin, Ethereum, etc. [16] Here, we have used one of the most popular consensus algorithms - Proof of Work (PoW) [31]. This ensures the security of the network in the form of block mining and the validity of the newly mined block [32]. In our experiment, each block in the network consists of a block header of key parameters, including block creation time, data from the input fasta file, associated header information, and reference to the previous block. Every time we want to create a new block in a network there is a need to check the level of difficulties measured by the PoW algorithm. It uses a block reference which must not exceed a certain threshold for the block to be considered valid. The computational power (or hash power) on the network is defined such that all possible variables in the block header are repeatedly iterated to find the value of B satisfying

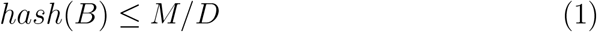

where D ∈[1, M] is the target difficulty. All possible variables in the block header are repeatedly iterated to find the value of B satisfying equation 1. The value of D is directly proportional to the number of iterations needed to find a valid block. The expected number of operations is exactly D [33].

### 3.2. Implementing Blockchain Model

Here we have proposed a general workflow diagram (figure 2) for implementing the Block Chain Model while accessing data for NGS data analysis. It encompasses blocks or nodes, which are connected through a network. The nodes communicate through a secure channel and use the hash value as a key. In this figure, Block 1 serves as the online resource for the sequencing data needed for further analysis. Block 2 and 3 access reads from fasta files stored in block 1. As the fasta file contains sequence ID and sequences, we need to collect all the sequence reads only from the fasta file, which is done by these two blocks. Block 4 is responsible for assembling and storing the final set of transcripts in HDFS. Here, the transcripts are assembled from reads coming from blocks 2 and 3 (i.e. building the transcripts from the reads coming from blocks 2 and 3 and storing the output in HDFS). We have used Ethereum techniques [34] for building blockchain networks. Ethereum is a global, public, and blockchain-based open-source platform for decentralized applications [35]. A block in an Ethereum blockchain is identified by a hash value of its content and references to the hash of the parent block. Hashes of blocks are created using cryptographic hash functions, which are mathematical algorithms that map data of arbitrary size to a bit string of a fixed size (called hash or digest). We have used R package *digest* [36, 37] for creating block and hash functions. Hash alone is not enough to prevent tampering. Here PoW algorithm has been used to solve this problem. It prevents denial of service attacks (DDoS) on a network and maintains privacy. We have assigned a hash value to each data packet (downloaded from NCBI) using the above-mentioned technique to maintain its privacy in the network. Subsequently, each data packet has been used further for transcript assembly.

**Figure 2.**
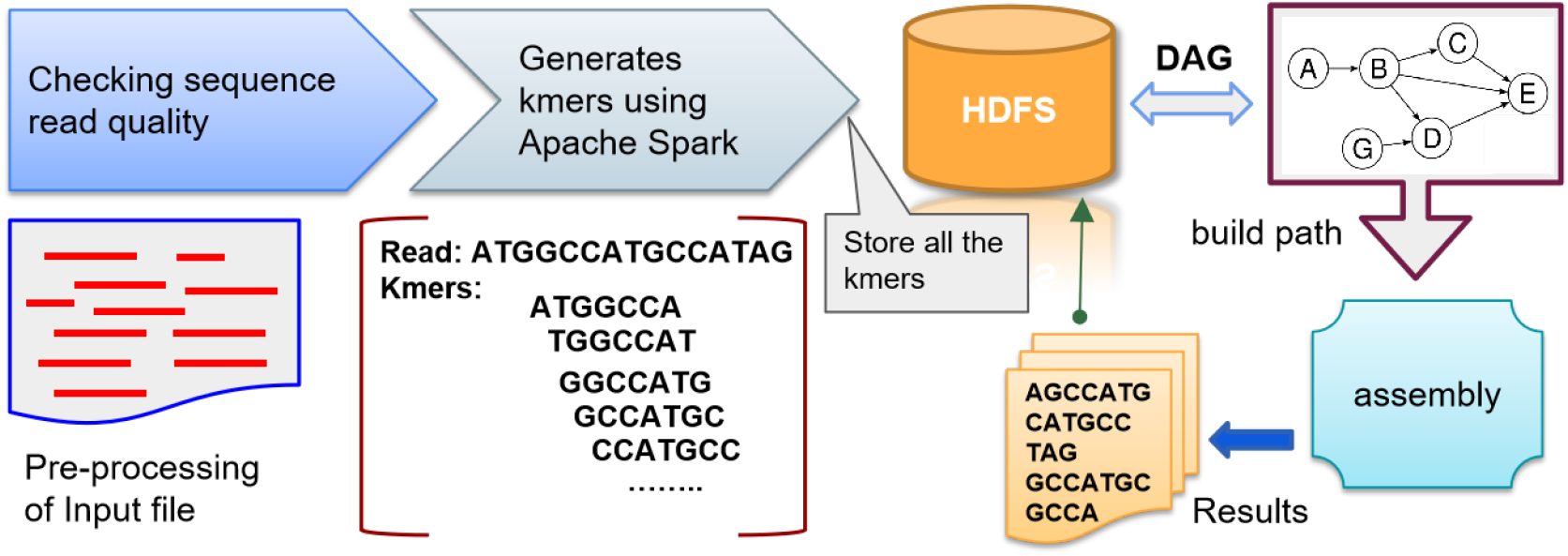
Workflow diagram for de novo transcript assembly

### 3.3. De novo transcript assembly

It consists of several steps including kmer generation, construction of graphs, and then assembly. These steps are described below and shown in figure 2.

#### 3.3.1. Creating k-mers and Construction of transcripts using de Bruijn graph

Most of the de novo assembly algorithms for short-read RNA sequences rely on the concept of the de Bruijn graph. In this paper, we have implemented de Bruijn graphs in two steps. In the first step, we have generated k-mers and in the second step, we have constructed the graph. The generation of all possible k-mers from the input sequences requires huge computation which makes it very time-consuming. Several existing software is available which generate k-mer but very few are distributed in nature. Here, we have used a Hadoop-based k-mer algorithm [38], developed in our lab, to reduce the execution time as it runs parallel on a distributed platform. We have further reduced the time using Apache Spark which allows in-memory computation. In algorithm 1, we have discussed the steps to generate a list of large amounts of k-mers and store it in HDFS or disk. In algorithm 1 step 2, we have also discussed the steps to construct the graph using all k-mer’s i.e. list of all k-mer stored or generated from step 1 and also generate all possible maximal simple paths.

##### Algorithm 1

Creating k-mers and Construction of transcripts

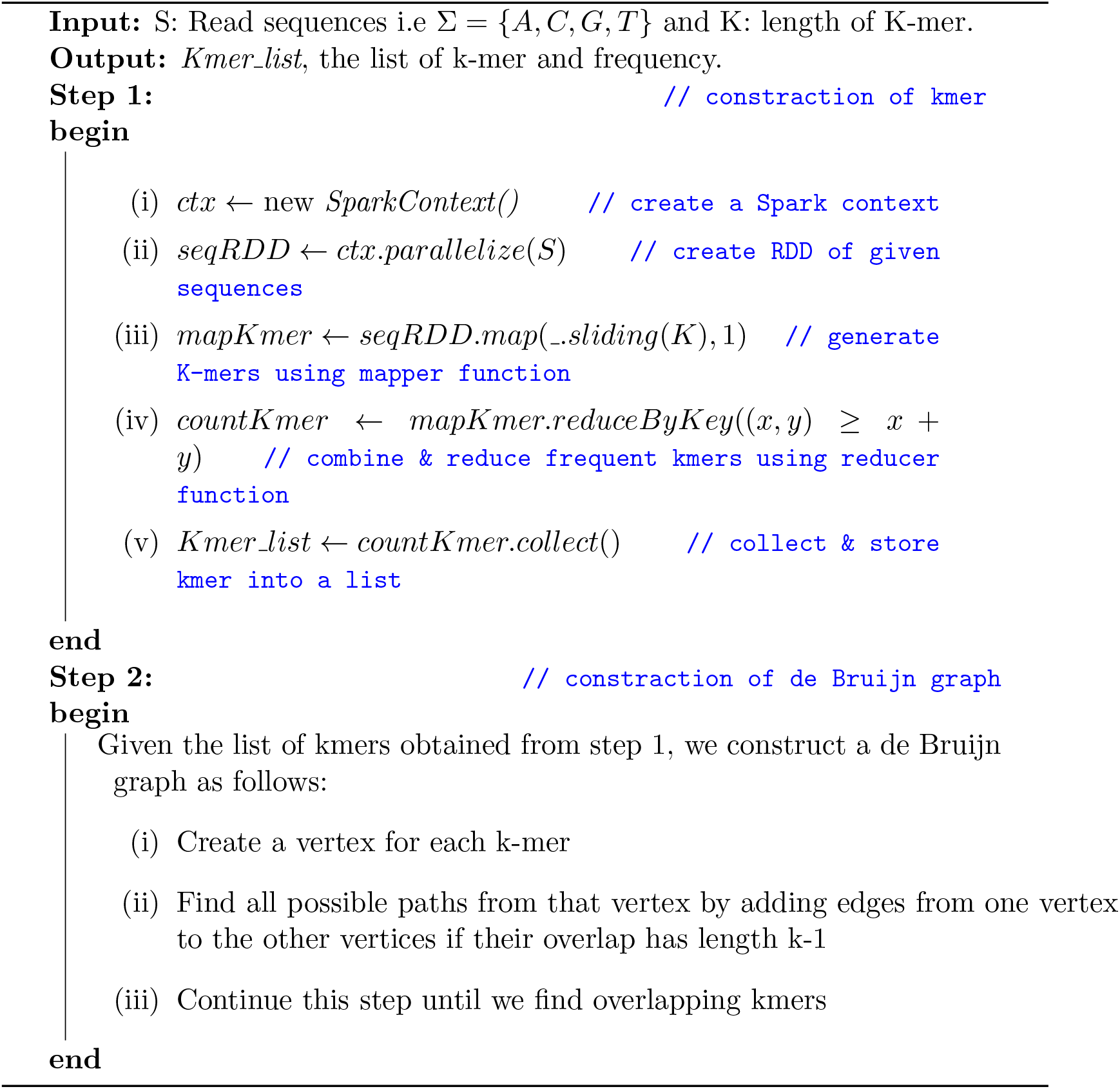

In algorithm 1, we have taken the Fasta sequence file that contains reads and the length of k-mer i.e. K as an input. We have used Apache Spark and created SparkContext [39] as a master Spark application. The sparkContext object allows the application to access the cluster for creating a resilient distributed dataset (RDD) using the parallelize() function. Each dataset (i.e. reads) in RDD is divided into many logical parts and it is computed on different nodes of the cluster. The flatMap() function is used to extract information from RDD. It operates on each row and returns multiple new rows for a given row. It parses the sequences from RDD to generate k-mer using a sliding function with fixed window-size K and sliding interval 1. It will create subsequences that overlap in k-1 positions. After processing, the mapper emits a *<*key,value*>* pair to the reducer where the key is a k-mer and the value is 1. The reduceByKey function collects each key and merges all the values emitted against the same key, i.e., *<*key, list(value)*>* pairs emitted by *flatmap()* function. In this function, (x, y) *≥* (x + y) initializes the x variable with default integer value 0 and adds up an element for each key. It combines the value of each identical k-mer and returns the final RDD with total counts paired with the key. i.e. if two keys are the same k-mer string then the value will be added until the list is empty. The reducer emits a final *<*key,value*>* pair where the key is a unique k-mer and the value is its frequency. The collect function is used to return the elements of the RDD generated by the reducer back to the driver program.

#### 3.3.2. Construction and optimization of transcripts length by SecDATA

Here, we have determined the optimal length of each transcript. The “Optimized length” represents the minimum number of nodes traversed for building all transcripts i.e. minimum path length for transcript construction. To do so, we have developed a directed acyclic graph (DAG) (described in algorithm 2) based method - GTraversal (described in algorithm 3).

In this paper, we have used DAG for de novo transcript reconstruction and quantification. We have used overlaps in k-mers to determine which k-mer pairs are adjacent in the read sequences. For example, if two k-mers <u>**TAGCAT**</u> and <u>**AGCATA**</u> overlap then they are adjacent nodes in the graph and there is a directed edge from <u>**TAGCAT**</u> to <u>**AGCATA**</u>. One k-mer may overlap over multiple other k-mers. Here we require computationally (N-1)(N)/2 steps for traversing the edges in the graph to find the possible directed paths (i.e. transcripts). A naive approach is to calculate the longest path from every node using the Depth First Search (DFS) algorithm. In our algorithm, we have used an adaptive DFS to traverse the DAG and also optimized the parameters for edge selection and path construction.

In algorithm 2, we have generated the DAG using an Adjacency List representation. This is a dictionary of *<*key,value*>* pairs, whose keys are the nodes of the graph. For each key, the corresponding value is a list containing the nodes that are adjacent to this key node. We have taken the k-mer generated from the previous step (discuss section 3.3.1) as input and stored it into a list called *kmer_list*. Algorithm 2 returns a dictionary that maps each k-mer in the *kmer_list* onto the set of all other k-mers in the list to which it overlaps with a minimal length min-overlap. Let *kmer*_1_ = *xy* and *kmer*_2_ = *yz* be two k-mers and *min-overlap = m* where m *∈* integers, then *suffix = y* with respect to *kmer*_1_ and *prefix=y* with respect to *kmer*_2_. One *<*key,value*>* pair of the dictionary, *<kmer*_1_, *kmer*_2_*>*, thus represents a directed edge that starts at a particular k-mer (here *kmer*_1_, used as the key in the dictionary) and lead to all other k-mers (here *kmer*_2_, used as the value in the dictionary) that overlap with it with of a minimal length m i.e. |*y*| *≥ m*. By definition of DAG, a k-mer, say *kmer*_1_, never points to itself, i,e. <key,value> ≠<*kmer*_1_, *kmer*_1_>, even though the k-mer might overlap itself and *<kmer*_1_, *kmer*_2_ *>* and *<kmer*_2_, *kmer*_1_*>* cannot co-exist in the dictionary as the graph is acyclic.

In **algorithm 3**, we have traversed the graph obtained in algorithm 2 to reassemble the kmers to build transcripts using adaptive depth-first search (DFS). DFS can be used to find all possible combinations of paths from a start vertex to all reachable vertices in the graph such that the ordering respects reachability. That is, if a vertex u is reachable from v, then v must be lower in the ordering, thus u must overlap on v.

We have used a recursive function, *find transript()*, to determine the path from a source kmer. It returns a list of nodes comprising the path that satisfies a minimum path length minlen. It takes the graph, start vertex, and path as arguments. Initially, the path is an empty list i.e. no nodes have been traversed yet. If the graph has the start vertex then we store it in the path. We recursively call *find transcript()* with the next node along with the current path that has already been traversed till it encounters untraversed reachable nodes. The same node will not occur more than once on the path returned i.e. it won’t contain cycles. We repeat this step for each kmer from the kmer-list.

In our experiment, we have considered the different combinations of overlap and minlength in algorithm 1 and algorithm 2 respectively. For example, let us consider five kmers generated from input reads (i.e. fasta file) (k1: TCGGA), (k2: GATTA), (k3: TTACA), (k4: ACAGA) and (k5: GGAAT). If we consider *overlap* = 2 and *minlen* = 2, the possible Adjacency list is *k*1 *→*[*k*2, *k*5], *k*2 *→*[*k*3], *k*3*→* [*k*4], *k*4*→* [*k*2] the path is [*k*1*− k*2*− k*3 *− k*4] and the transcript is “**TCGGATTACAGA**”. But if we consider *overlap* = 3 and *minlen* = 2, the possible Adjacency list is {*k*1 *→* [*k*5], *k*2*→* [*k*3], *k*3 *→* [*k*4] } and the path is [*k*2 *−k*3 *−k*4] and the transcript is “**GATTACAGA**”.

##### Algorithm 2

**CreateDAG()**

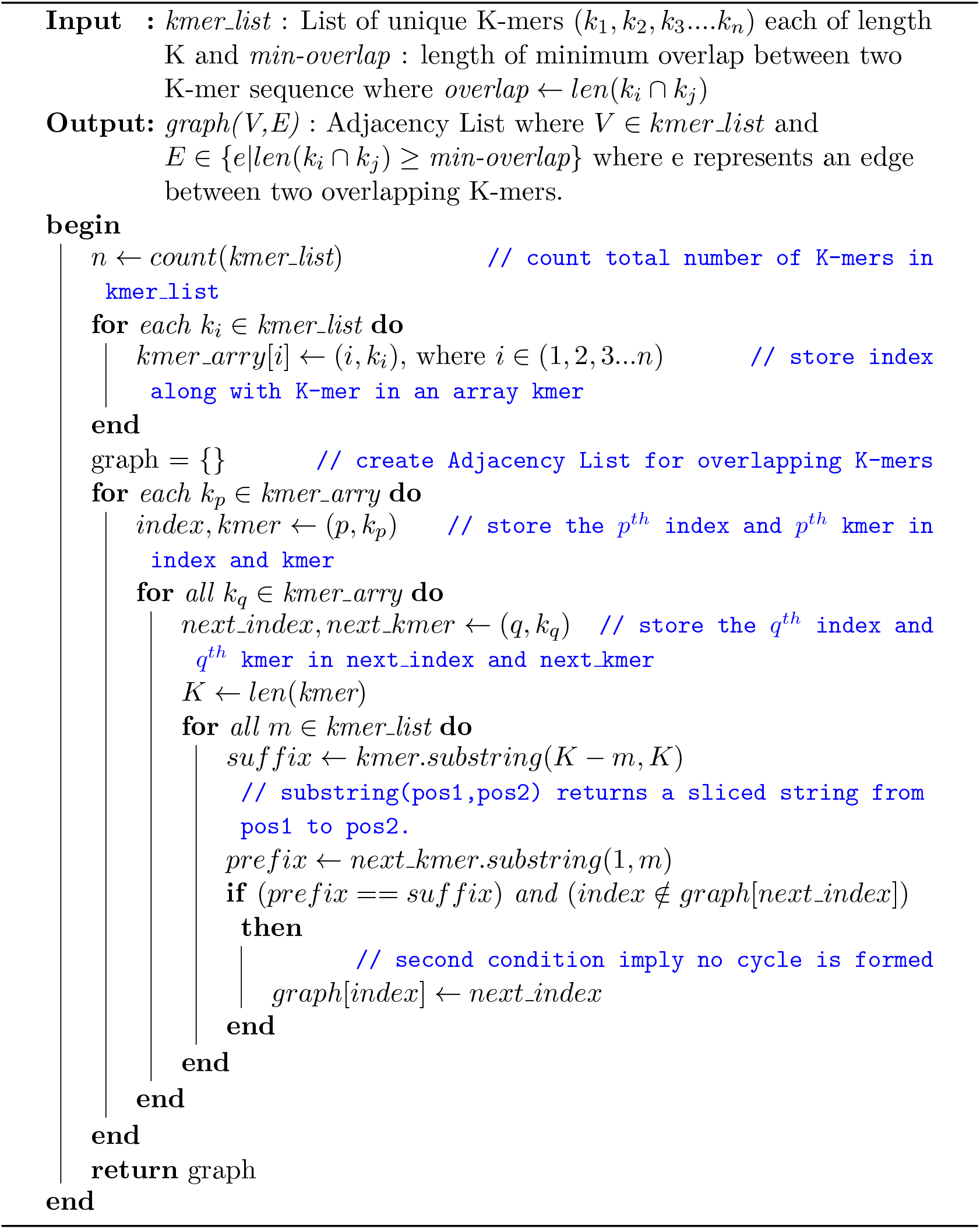

##### Algorithm 3

**GTraversal()**

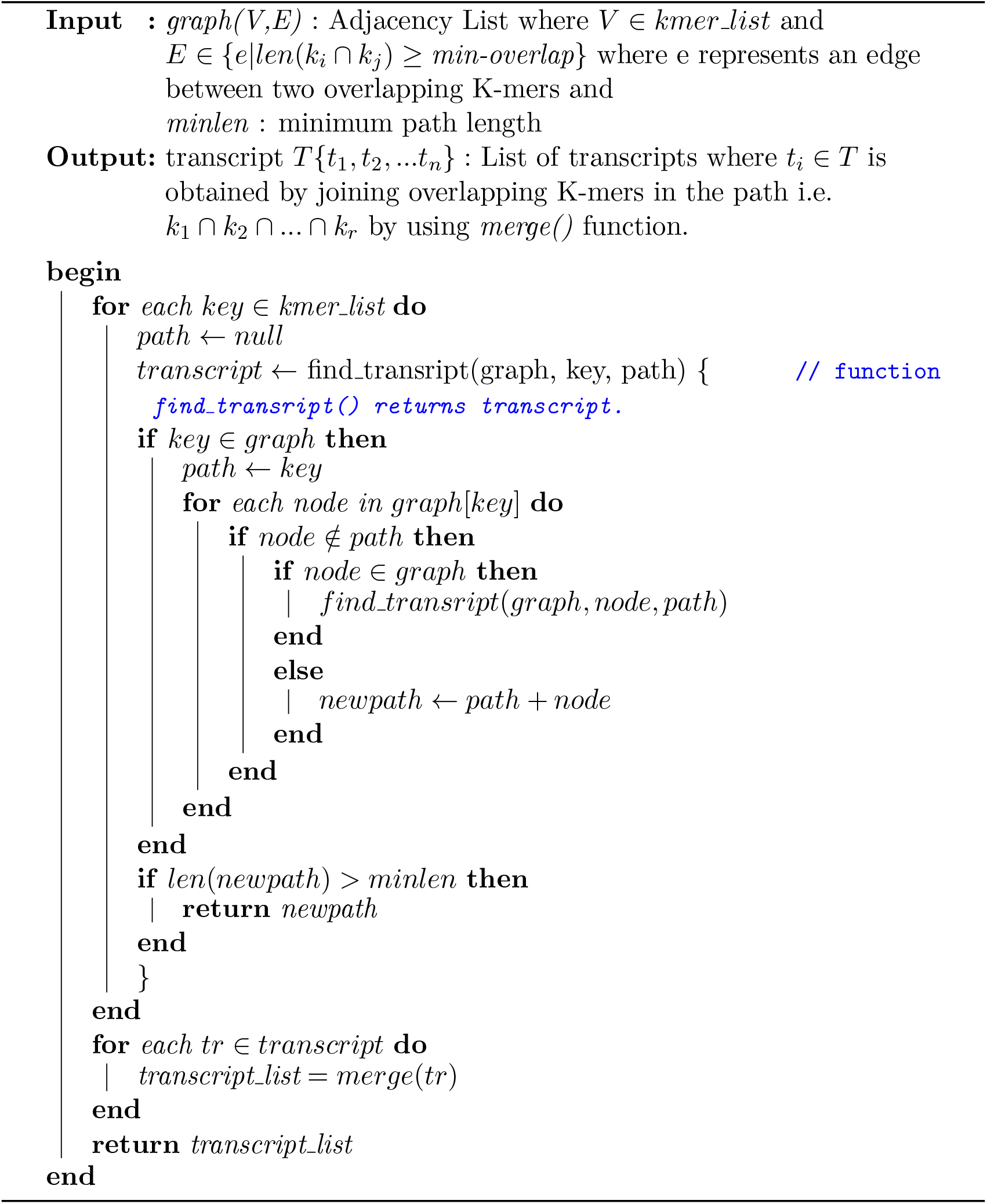

We have summarized the final optimized length of each input dataset including simulated reads. Figure 3 summarizes the optimized length thus obtained, where the y-axis represents the total number of reads in input files, the x-axis represents the input files, and the final optimized transcript lengths for each input file are demarcated in the figure.

**Figure 3.**
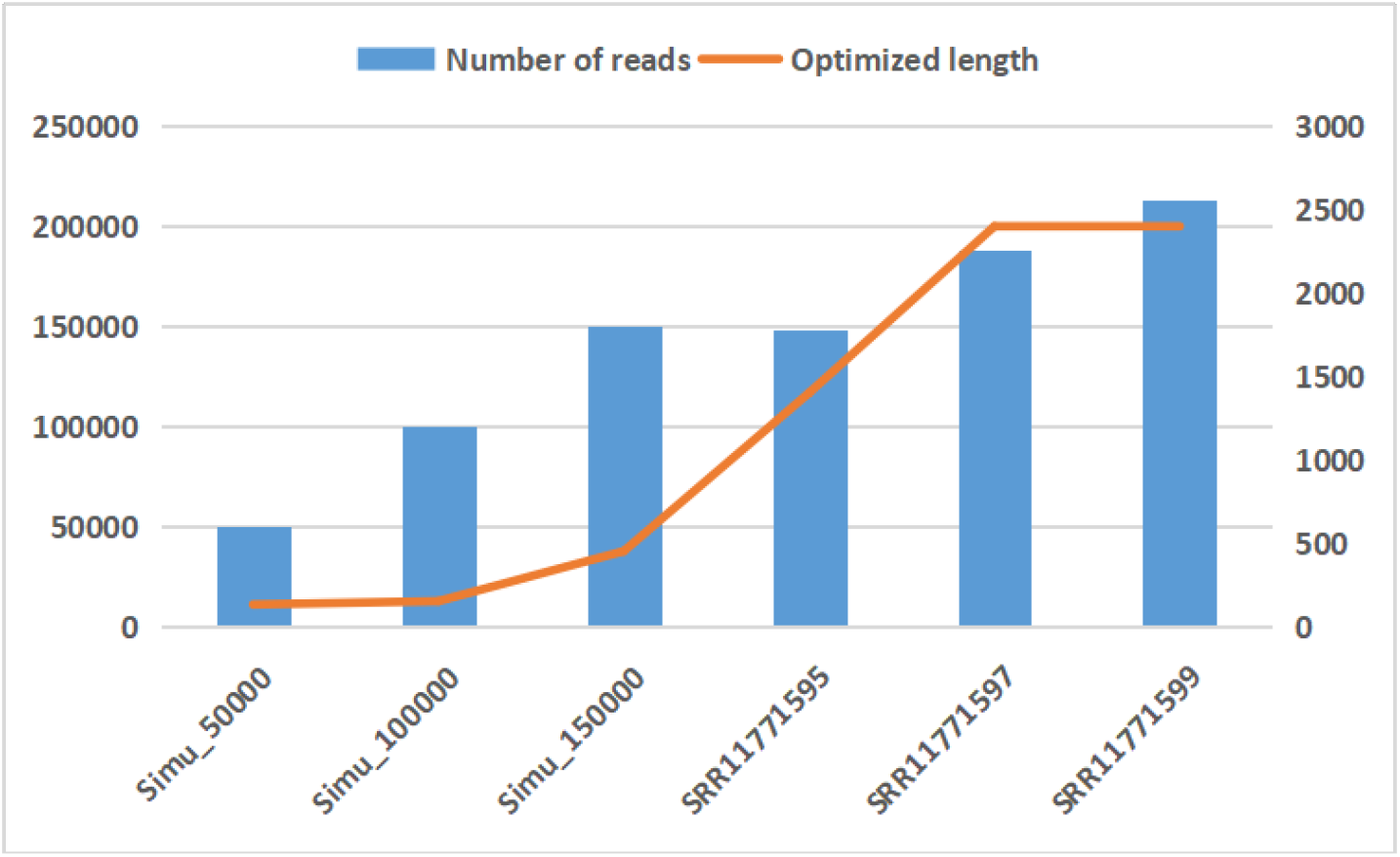
Computational framework for SecDATA

## 4. Results and Discussion

In this experiment, we have taken both simulated (discussed in section 2.1) and real online human datasets (as shown in Table 1) for our analysis. We have evaluated the performance on machines with an i7 processor CPU 3.6 GHz x 8 with 8GB of RAM.

### 4.1. Mining difficulty

The difficulty[40] indicates how difficult it is to find the hash of a new block. We have ensured the security of the accessed sequence data by measuring the difficulty level of creating a new block (discussed in section 3.2). Here, the ‘difficulty’ level explains, how hard it is to add a fake block to the blockchain network involving some attempts for a particular period. Figure 4 shows the difficulty level along the X-axis concerning the number of iterations, along the Y-axis using the Pow algorithm. Iterated Logarithm or Log*(n), representing the number of attempts to add a new block, is the number of times the logarithm function must be iteratively applied till the result *≤* 1. A high difficulty indicates that mining the same number of blocks will need more computational power, making the network more secure against threats. Figure 4 shows that increasing the difficulty level causes the iteration to rise on a specific part before converging.

**Figure 4.**
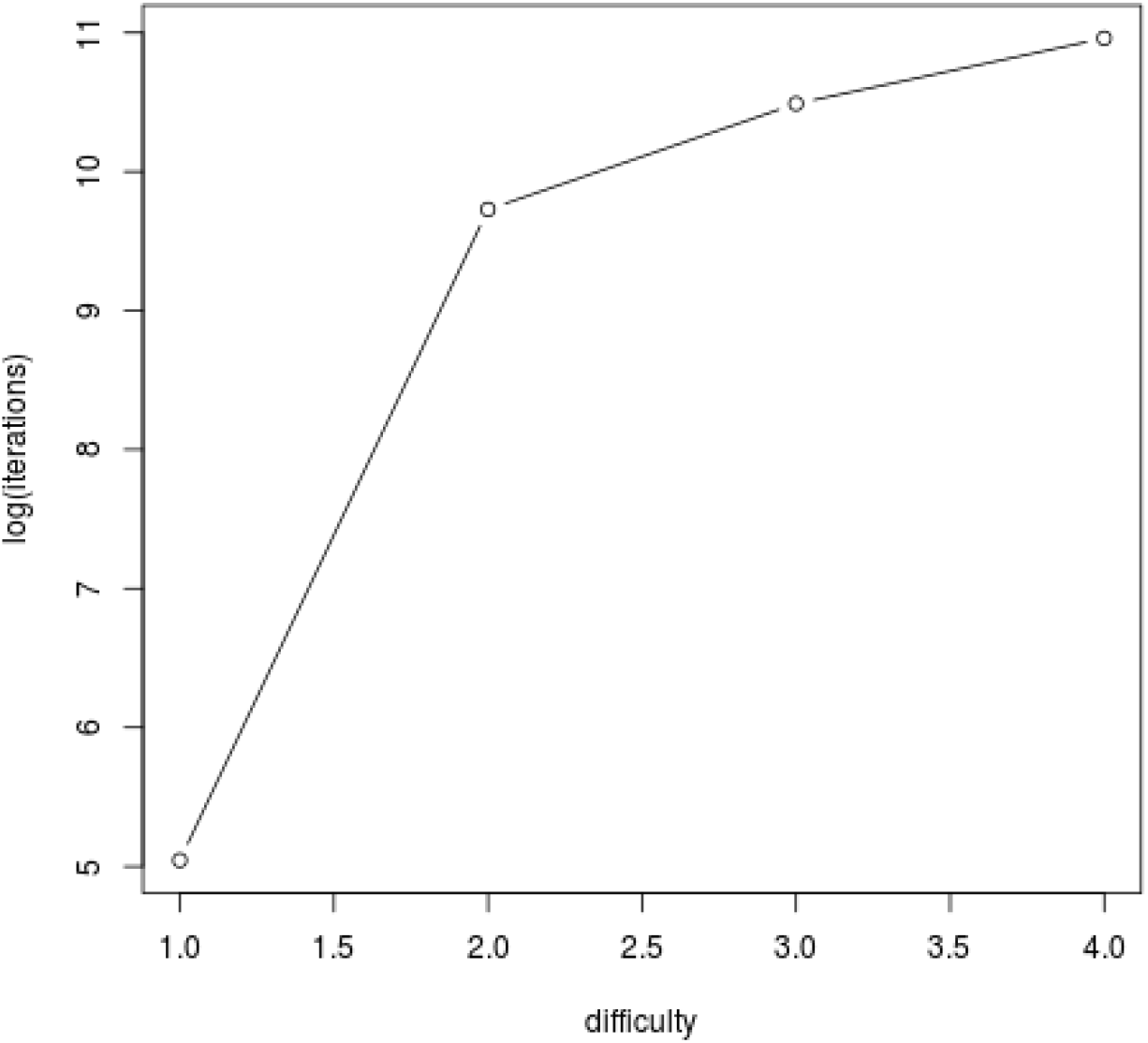
The difficulty level of creating new block with respect to iteration using Pow algorithm

### 4.2. Creating k-mers using different computational platforms

As discussed in section 3.3.1, k-mer is an important but time-consuming process for de novo transcript assembly. Further, this is the initial step for transcript assembly. Hence, we have compared the execution time for generating k-mer in different computational platforms including stand-alone and distributed framework (which includes Spark and Hadoop). In the case of Hadoop, we observed the requirement of much less execution time for generating k-mers(as shown in figure 5). To reduce the time further, we have used the Spark-based method and have compared this result with that from Hadoop shown in figure 5.

**Figure 5.**
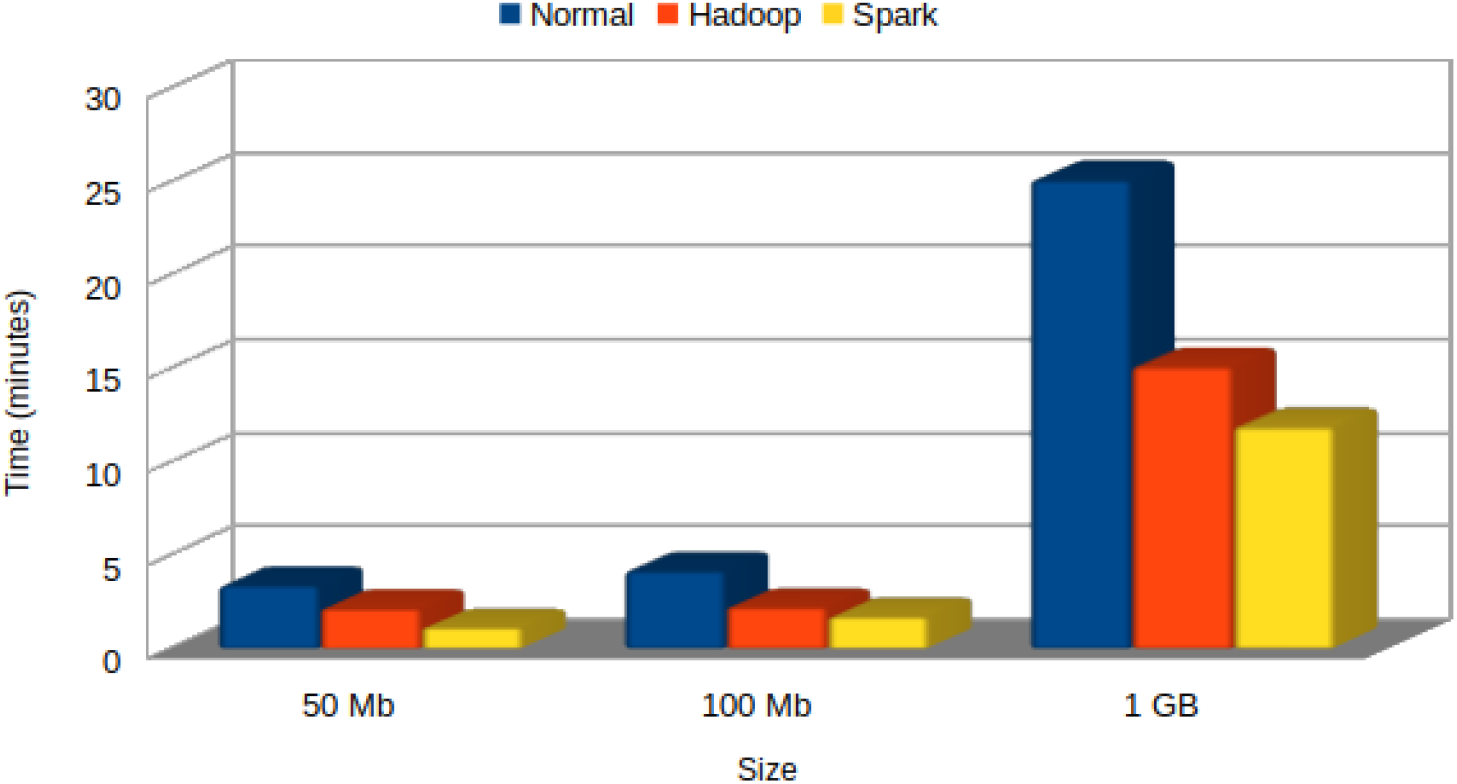
Time comparison of generating kmer using standalone, Hadoop, and spark framework

Here, the x-axis represents the size of the data, and the y-axis represents the execution time in minutes. Three differently sized datasets have been used as input for checking the execution time in various computational platforms. We have considered the kmer length of 21 i.e. k=21 for all the experiments in this work. In comparison to standalone and Hadoop, Apache Spark outperforms with respect to execution time.

### 4.3. Evaluating the performance of SecDATA

Transcript Assembly: To evaluate the performance of SecDATA, we have compared its efficiency with that of Debruijn and Trinity. The efficiency has been calculated (in terms of alignment percentage) based on the difference between expected and actual output. To generate the expected output for all these protocols, we have performed the following:

#### 4.3.1. Sequence Alignment using BWA

We aligned the resultant transcripts from Debruijn, Trinity, and SecDATA using the local alignment tool BWA [41] with human (hg18) and mouse (mm18) reference genome. Here we have performed a comparison of the alignment score for both sets of input (table 1). In the case of all the simulated datasets, we observed (in figure 6b) the maximum alignment score for the transcripts obtained by using SecDATA and Trinity. On the contrary, for transcripts obtained by using de Bruijn, the alignment percentage is 21.22%, 17.13%, and 16.7% respectively for simu 50000, simu 100000, and simu 150000. Figure 6a reveals a similar take-home message from the alignment results obtained by applying the 3 protocols on SRA datasets viz. SRR11771595, SRR11771595, SRR11771595 (figure 6a).

**Figure 6.**
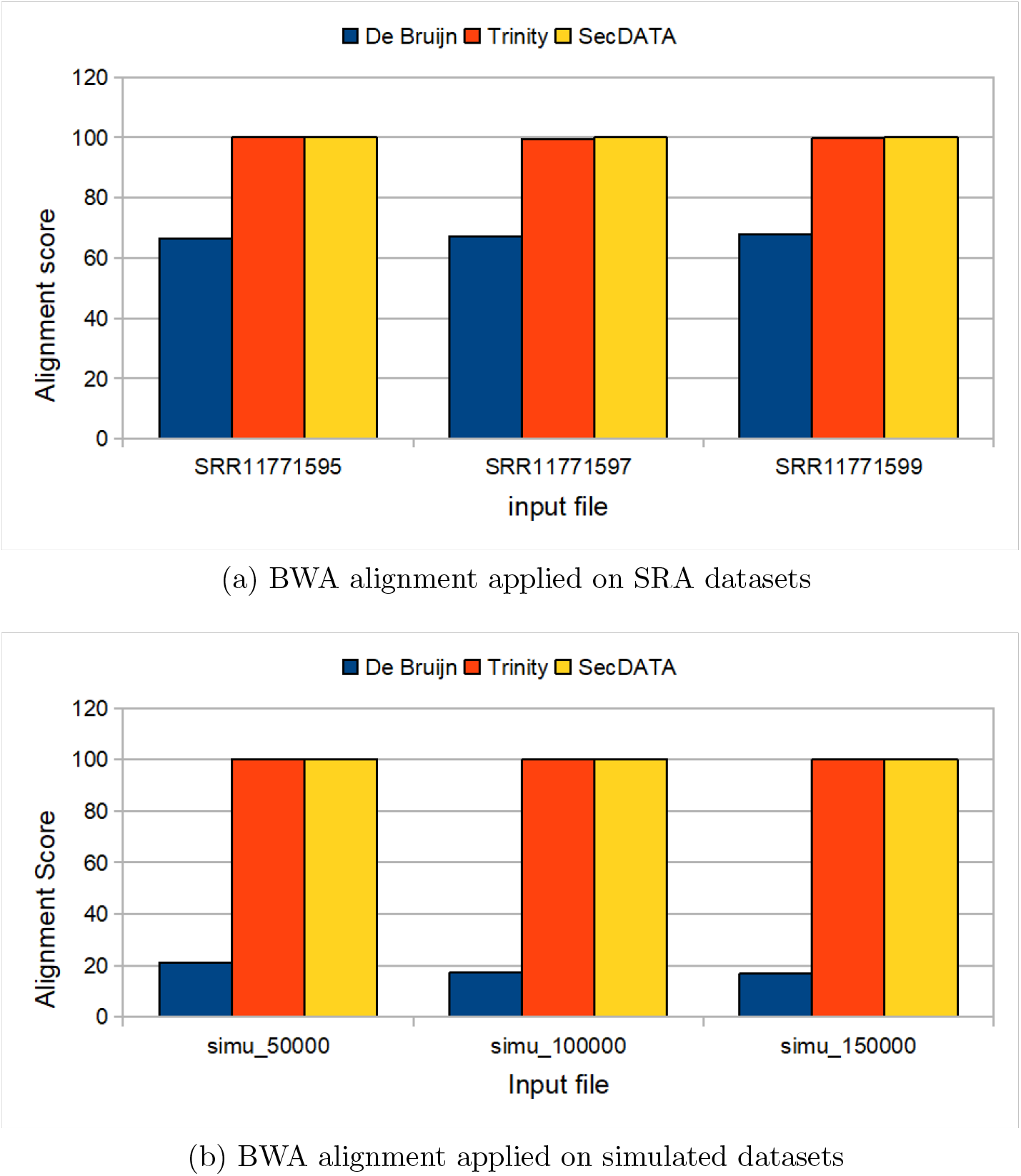
BWA alignment percentage obtained on outputs from de Bruijn, Trinity, and SecDATA

#### 4.3.2. Sequence Alignment using Blat

To check the efficiency of these three protocols for the number of true positive outputs, we have compared our results with the output of the reference-based transcript assembly tool using the Blat similarity search tool [42]. In short, we have obtained Human hg38 and Mouse mm10 reference genome and annotation (GTF) files from Gencode (www.gencodegenes.org). This was followed by genome alignment and transcript assembly of the corresponding Human and Mouse SRA and simulated files with Hisat2[43] and Stringtie[43] tools respectively. Considering these detected transcript sequences as a reference, BLAT was run with the output of de novo assembled sequences. Figure 7 shows the Blat results (expressed in terms of alignment percentage based on the difference between expected and actual output) obtained on outputs from de Bruijn, Trinity and SecDATA applied on both the SRA (Figure 7a) and simulated datasets (Figure 7b). We have considered no gap here i.e. each transcript matches without any gap to the reference transcript file. At this stage, a significant difference is observed in the alignment percentage obtained on outputs from all the 3 protocols. SecDATA outperforms both Trinity and de Bruijn.

**Figure 7.**
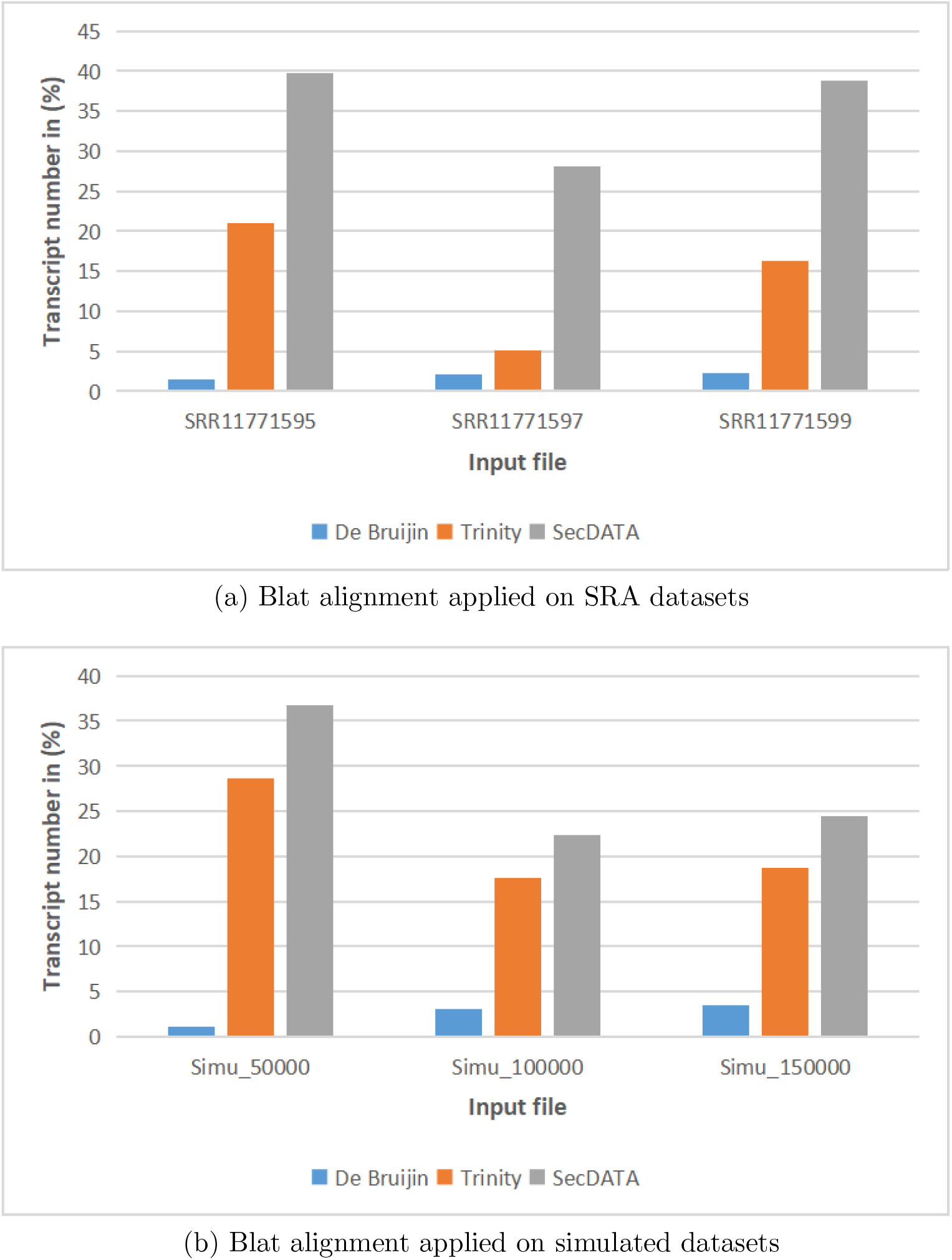
Blat alignment result obtained on outputs of de Bruijn, Trinity, and SecDATA

## Conclusion

In this paper, we presented a novel protocol *SecDATA* for de novo transcript assembly from RNA-seq data. The novelty of this work relies on (i) secured input data access and storage of the output and (ii) efficient de novo transcript assembly using optimized DAG. Here, we have introduced a blockchain-based secure data access and storage technique that mitigates the reliability issue of the input dataset to be used for transcript assembly. For data security, we have used a POW algorithm to verify the creation of a false block in a network. For de novo transcript assembly, we generated the k-mers using spark and then created as well as traversed the DAG to build the transcripts. Here, we have used an adaptive DFS to traverse the DAG. Further, we have determined the optimal length of each transcript.

Our protocol outperforms other state-of-the-art transcript assembly protocols -Trinity and de Bruijn in terms of the output i.e. number of true positive transcripts. We plan to implement *SecDATA* in a cloud environment so that it can be accessed as a service in the future.

SecDATA is available at https://github.com/sudipmondalcse/SecDATA.

## Conflicts of Interest

The authors declare that they have no conflicts to disclose.

## Notes

### Competing Interest Statement

The authors have declared no competing interest.

## References

[1] K.-O. Mutz, A. Heilkenbrinker, M. Lönne, J.-G. Walter, F. Stahl, Transcriptome analysis using next-generation sequencing, Current opinion in biotechnology 24 (1) (2013) 22–30.

[2] M. Geniza, P. Jaiswal, Tools for building de novo transcriptome assembly, Current Plant Biology 11 (2017) 41–45.

[3] M.-X. Chen, F.-Y. Zhu, B. Gao, K.-L. Ma, Y. Zhang, A. R. Fernie, Chen, L. Dai, N.-H. Ye, X. Zhang, et al., Full-length transcript-based proteogenomics of rice improves its genome and proteome annotation, Plant Physiology 182 (3) (2020) 1510–1526.

[4] E. Bushmanova, D. Antipov, A. Lapidus, A. D. Prjibelski, rnaspades: a de novo transcriptome assembler and its application to rna-seq data, GigaScience 8 (9) (2019) giz100.

[5] J. A. Martin, Z. Wang, Next-generation transcriptome assembly, Nature Reviews Genetics 12 (10) (2011) 671–682.

[6] Y. Peng, H. C. Leung, S.-M. Yiu, M.-J. Lv, X.-G. Zhu, F. Y. Chin, Idba-tran: a more robust de novo de bruijn graph assembler for transcriptomes with uneven expression levels, Bioinformatics 29 (13) (2013) i326–i334.

[7] M. Hölzer, M. Marz, De novo transcriptome assembly: A comprehensive cross-species comparison of short-read rna-seq assemblers, Gigascience 8 (5) (2019) giz039.

[8] Z. Li, Y. Chen, D. Mu, J. Yuan, Y. Shi, H. Zhang, J. Gan, N. Li, Hu, B. Liu, et al., Comparison of the two major classes of assembly algorithms: overlap–layout–consensus and de-bruijn-graph, Briefings in functional genomics 11 (1) (2012) 25–37.

[9] K. Clarke, Y. Yang, R. Marsh, L. Xie, et al., Comparative analysis of de novo transcriptome assembly, Science China Life Sciences 56 (2) (2013) 156–162.

[10] R. Rizzi, S. Beretta, M. Patterson, Y. Pirola, M. Previtali, G. Della Vedova, P. Bonizzoni, Overlap graphs and de bruijn graphs: data structures for de novo genome assembly in the big data era, Quantitative Biology 7 (4) (2019) 278–292.

[11] P. E. Compeau, P. A. Pevzner, G. Tesler, How to apply de bruijn graphs to genome assembly, Nature biotechnology 29 (11) (2011) 987–991.

[12] R. Chikhi, A. Limasset, S. Jackman, J. T. Simpson, P. Medvedev, On the representation of de bruijn graphs, in: International conference on Research in computational molecular biology, Springer, 2014, pp. 35–55.

[13] A. Bowe, T. Onodera, K. Sadakane, T. Shibuya, Succinct de bruijn graphs, in: International workshop on algorithms in bioinformatics, Springer, 2012, pp. 225–235.

[14] H. I. Ozercan, A. M. Ileri, E. Ayday, C. Alkan, Realizing the potential of blockchain technologies in genomics, Genome research 28 (9) (2018) 1255–1263.

[15] H. Natarajan, S. Krause, H. Gradstein, Distributed ledger technology and blockchain (2017).

[16] A. Sunyaev, Distributed ledger technology, in: Internet Computing, Springer, 2020, pp. 265–299.

[17] G. Fiorentino, C. Occhipinti, A. Corsi, E. Moro, J. Davies, A. Duke, Blockchain: Enabling trust on the internet of things, The Internet of Things: From Data to Insight (2020) 141–157.

[18] B. Farahani, F. Firouzi, M. Luecking, The convergence of iot and distributed ledger technologies (dlt): Opportunities, challenges, and solutions, Journal of Network and Computer Applications 177 (2021) 102936.

[19] K. Jyothilakshmi, V. Robins, A. Mahesh, A comparative analysis between hyperledger fabric and ethereum in medical sector: A systematic review, Sustainable Communication Networks and Application (2022) 67–86.

[20] M. Valenta, P. Sandner, Comparison of ethereum, hyperledger fabric and corda, Frankfurt School Blockchain Center 8 (2017) 1–8.

[21] S. Thiebes, M. Schlesner, B. Brors, A. Sunyaev, Distributed ledger technology in genomics: a call for europe, European Journal of Human Genetics 28 (2) (2020) 139–140.

[22] S. Mondal, R. K. Maji, Z. Ghosh, S. Khatua, Parstream-seq: An improved method of handling next generation sequence data, Genomics 111 (6) (2019) 1641–1650.

[23] A. M. Bolger, M. Lohse, B. Usadel, Trimmomatic: a flexible trimmer for illumina sequence data, Bioinformatics 30 (15) (2014) 2114–2120.

[24] J. Brown, M. Pirrung, L. A. McCue, Fqc dashboard: integrates fastqc results into a web-based, interactive, and extensible fastq quality control tool, Bioinformatics 33 (19) (2017) 3137–3139.

[25] H. Gourlé, O. Karlsson-Lindsjö, J. Hayer, E. Bongcam-Rudloff, Simulating illumina metagenomic data with insilicoseq, Bioinformatics 35 (3) (2019) 521–522.

[26] N. Z. Benisi, M. Aminian, B. Javadi, Blockchain-based decentralized storage networks: A survey, Journal of Network and Computer Applications 162 (2020) 102656.

[27] K. Wüst, A. Gervais, Do you need a blockchain?, in: 2018 Crypto Valley Conference on Blockchain Technology (CVCBT), IEEE, 2018, pp. 45–54.

[28] K. Sultan, U. Ruhi, R. Lakhani, Conceptualizing blockchains: Characteristics & applications, arXiv preprint arXiv:1806.03693 (2018).

[29] T. M. Fernández-Caramés, P. Fraga-Lamas, Advances in the convergence of blockchain and artificial intelligence (2022).

[30] D. Mingxiao, M. Xiaofeng, Z. Zhe, W. Xiangwei, C. Qijun, A review on consensus algorithm of blockchain, in: 2017 IEEE international conference on systems, man, and cybernetics (SMC), IEEE, 2017, pp. 2567–2572.

[31] L. M. Bach, B. Mihaljevic, M. Zagar, Comparative analysis of blockchain consensus algorithms, in: 2018 41st International Convention on Information and Communication Technology, Electronics and Microelectronics (MIPRO), Ieee, 2018, pp. 1545–1550.

[32] T. Xue, Y. Yuan, Z. Ahmed, K. Moniz, G. Cao, C. Wang, Proof of contribution: A modification of proof of work to increase mining efficiency, in: 2018 IEEE 42nd annual computer software and applications conference (COMPSAC), Vol. 1, IEEE, 2018, pp. 636–644.

[33] G. BitFury, Proof of stake versus proof of work, White paper, Sep (2015).

[34] A. E. Gencer, S. Basu, I. Eyal, R. v. Renesse, E. G. Sirer, Decentralization in bitcoin and ethereum networks, in: International Conference on Financial Cryptography and Data Security, Springer, 2018, pp. 439–457.

[35] C. BouSaba, E. Anderson, Degree validation application using solidity and ethereum blockchain, in: 2019 SoutheastCon, IEEE, 2019, pp. 1–5.

[36] http://dirk.eddelbuettel.com/code/digest.html (2020).

[37] D. Eddelbuettel, A. Lucas, J. Tuszynski, H. Bengtsson, S. Urbanek, M. Frasca, B. Lewis, M. Stokely, H. Muehleisen, D. Murdoch, et al., Package ‘digest’ (2022).

[38] S. Mondal, S. Khatua, Finding simple sequence repeats (ssrs) within human genome using mapreduce based k-mer algorithm, in: 2018 Fifth International Conference on Parallel, Distributed and Grid Computing (PDGC), IEEE, 2018, pp. 340–345.

[39] S. Chellappan, D. Ganesan, Introduction to apache spark and spark core, in: Practical Apache Spark, Springer, 2018, pp. 79–113.

[40] G. A. Pierro, H. Rocha, The influence factors on ethereum transaction fees, in: 2019 IEEE/ACM 2nd International Workshop on Emerging Trends in Software Engineering for Blockchain (WETSEB), IEEE, 2019, pp. 24–31.

[41] H. Li, R. Durbin, Fast and accurate long-read alignment with burrows– wheeler transform, Bioinformatics 26 (5) (2010) 589–595.

[42] W. J. Kent, Blat—the blast-like alignment tool, Genome research 12 (4) (2002) 656–664.

[43] M. Pertea, D. Kim, G. M. Pertea, J. T. Leek, S. L. Salzberg, Transcript-level expression analysis of rna-seq experiments with hisat, stringtie and ballgown, Nature protocols 11 (9) (2016) 1650–1667.

